# GLiDe: a web-based genome-scale CRISPRi sgRNA design tool for prokaryotes

**DOI:** 10.1101/2022.11.25.517898

**Authors:** Tongjun Xiang, Huibao Feng, Xin-hui Xing, Chong Zhang

**Affiliations:** MOE Key Laboratory for Industrial Biocatalysis, Institute of Biochemical Engineering, Department of Chemical Engineering, Tsinghua University, Beijing 100084, China; Division of Biology and Bioengineering, California Institute of Technology, Pasadena, CA 91125, USA; Center for Synthetic and Systems Biology, Tsinghua University, Beijing 100084, China; Institute of Biopharmaceutical and Health Engineering, Tsinghua Shenzhen International Graduate School, Shenzhen 518055, China

## Abstract

CRISPRi screening has become a powerful approach for functional genomic research. However, the off-target effects resulting from the mismatch tolerance between sgRNAs and their intended targets is a primary concern in CRISPRi applications. To address this issue, we introduce Guide Library Designer (GLiDe), a web-based tool specifically created for the genome-scale design of sgRNA libraries tailored for CRISPRi screening in prokaryotic organisms. GLiDe incorporates a robust quality control framework, rooted in prior experimental knowledge, ensuring the accurate identification of off-target hits. It boasts an extensive built-in database, encompassing 1,397 common prokaryotic species as a comprehensive design resource. In addition, GLiDe provides the capability to design sgRNAs for newly discovered organisms. We further demonstrated that GLiDe exhibits enhanced precision in identifying off-target binding sites for the CRISPRi system.

**Availability and Implementation:** Freely available on the web at: https://www.thu-big.net/sgRNA_design/.

**AUTHOR SUMMARY:** CRISPRi screening is a powerful strategy for functional genomic study in prokaryotes. However, its applicability has been hindered by false positive results arising from off-target binding of sgRNAs. To tackle this challenge head-on, we present GLiDe, a web-based tool meticulously crafted for the genome-scale CRISPRi sgRNA library design in prokaryotes. GLiDe integrates a stringent quality control mechanism grounded in prior experimental insights. It has a built-in database and accepts uploaded reference files for new organisms. Moreover, we have demonstrated that our tool outperforms existing tools in CRISPRi sgRNA design for prokaryotes through cytometry assay. In summary, GLiDe emerges as a robust solution for sgRNA library design, poised to significantly advance functional genomic studies.

## INTRODUCTION

As the number of bacterial genomes continues to grow, genotype-phenotype mapping is emerging as a potent approach in functional genomics research. It yields valuable insights into the microbiology and engineering of microorganisms [1–3]. To investigate the function of a specific gene, the most straightforward strategy is to perturb the gene and then observe the resulting changes in phenotype [4]. In recent years, CRISPR system originates from bacterial immune system has provided an extraordinary tool for genome editing, which is programmable and applicable in a wide range of organisms [5–7]. By inactivating the endonuclease activity of Cas9 through mutations (D10A and H840A), nuclease-dead Cas9 (dCas9) is generated [8]. This can be adapted for transcriptional regulation, giving rise to the CRISPR interference (CRISPRi) system [9]. This innovative approach facilitates CRISPRi screening [10], a technique for genome-wide gene perturbation. This method employs a pooled library of single-guide RNA (sgRNA), where each sgRNA is meticulously designed to guide dCas9 protein to a specific genetic locus flanked by 3’-NGG protospacer adjacent motif (PAM) via Watson-Crick base pairing [8], resulting in the inhibition of transcription of the target gene. Following the introduction of the sgRNA library, high-throughput screening techniques such as fluorescence-activated cell sorting [11,12] or fitness screening [13,14] are used to assess how cells with different sgRNAs behave. These behaviors are subsequently identified through next-generation sequencing. However, false positive results may arise occasionally in CRISPRi screening, primarily due to unexpected binding, a phenomenon commonly known as “off-targeting.” This can occur due to the tolerance of mismatches between sgRNAs and their intended targets [15]. Consequently, off-target binding may lead to the unintended inhibition of additional genes beyond the originally targeted one. Hence, ensuring highly reliable sgRNA design is of paramount importance.

To date, a variety of tools have been established for sgRNA design [16–22]. It’s worth noting that these applications are predominantly tailored for the CRISPR/Cas9 system, where target DNA cleavage typically takes place following a conformational gating process [23]. This process requires sufficient base pairing between the sgRNA and the target DNA. In contrast, the CRISPRi system, which we are focusing on, involves inhibiting gene expression rather than cleaving DNA. In CRISPRi, off-target binding is more prone to occur compared to off-target cleavage [24], presenting challenges in accurately pinpointing off-target binding occurrences. Additionally, it’s crucial to consider the context of prokaryotic genomes when discussing sgRNA design tools. Prokaryotic genomes differ fundamentally from eukaryotic genomes in terms of genome accessibility due to a less organized chromosome structure [25,26] and variations in intracellular conditions. These unique characteristics may impact the applicability of existing sgRNA design tools to prokaryotic systems.

To tackle the challenges outlined above, our research focuses on the development of an accurate off-target identification algorithm tailored specifically for the CRISPRi system in prokaryotes. Drawing upon insights gained from extensive pooled screening assays evaluating binding affinities of diverse sequences in prokaryotes [10,27], we have conducted a comprehensive analysis to discern the impact of mismatches on binding affinity. From these derived principles, we have developed CRISPRi Guide Library Designer (GLiDe), a web-based tool that allows users to easily and rapidly design genome-scale sgRNA libraries for CRISPRi screening in prokaryotes. GLiDe offers a built-in database of 1,397 common microorganisms and accepts uploaded reference files for less common species. Additionally, we illustrated that sgRNAs generated by GLiDe display a reduced propensity for off-target binding. Our in-depth analysis, supported by quantitative data, unequivocally demonstrated that GLiDe showcased substantial performance improvement in designing sgRNA libraries for CRISPRi screening. Consequently, through rigorous design principles and meticulous experimentation, we have firmly established GLiDe as a highly promising tool poised to substantially promote functional genomic studies in prokaryotes.

## RESULTS

### Implementation of the GLiDe web tool

We have developed GLiDe, a web-based tool for genome-scale CRISPRi sgRNA library design in prokaryotes, which is available at https://www.thu-big.net/sgRNA_design/. It employs a powerful algorithm for identifying off-target hits and outputs the results on an interactive page (Figure 1A). This result can also be easily downloaded from the website in tabular format (Figure 1B, C).

**Figure 1.**
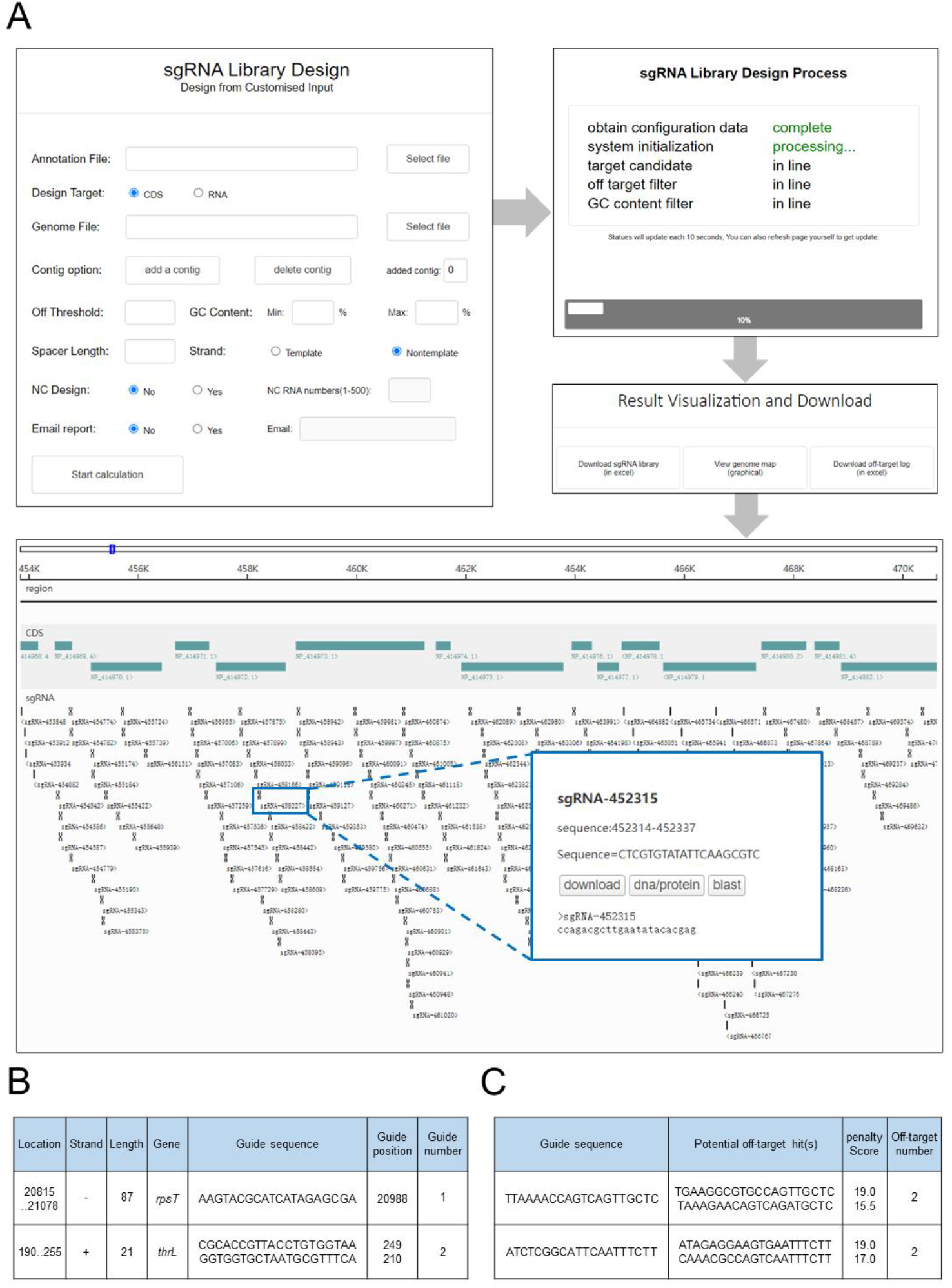
Workflow and results of a GLiDe query. **(A)** The main design process starts when a “start calculation” request is received, and the website automatically redirects to the progress page that displays the real-time progress of the calculation. After the calculation is complete, GLiDe redirects again to the results page. Users are free to download the result tables and view the sgRNA library on the interactive page. **(B)** The designed sgRNA library is grouped by its targeted gene. **(C)** The list of sgRNAs which are excluded from quality control, along with their off-target hits and penalty scores.

### Input and configuration

GLiDe includes an extensive built-in database harboring over 1,397 prokaryotic organisms and allows users to upload their reference files (see **Materials and Methods**). To optimize sgRNA design, five essential parameters are required: (i) design target, which can be either coding sequence (CDS) or RNA coding genes (RNA); (ii) off-target threshold, which is the penalty score used in the sgRNA quality control (see **Materials and Methods**); (iii) GC limits, which represent the upper and lower boundaries for the GC content of each sgRNA; (iv) spacer length, which defines the length of the sgRNA being designed; and (v) target strand, which can be either template or non-template, referring to the strand that the sgRNAs target. It’s important to note that, although for common CRISPRi systems, the non-template strand is the preferred choice to ensure effective gene silencing [9], we offer the alternative option for other applications, such as genome editing, base editing [28,29], or dynamic imaging [30], where there may be less strand preference.

GLiDe can also design library for multi-contig sequences, such as strains containing plasmids. For the second and subsequent contigs, the sequence file is necessary, while the annotation file is optional. This feature is designed for contigs where there is no need for sgRNA design, but the prevention of sgRNAs targeting these specific contigs is desired, like plasmid vectors used in experiments. If users choose to upload the annotation file for these additional contigs, GLiDe will design sgRNAs accordingly.

### Process and output

Upon configuration, a server request is initiated to commence the workflow (Figure 2). Anticipated processing time for a standard genome-scale library design range from 5 to 20 minutes, varying based on the size of the genome. The result sgRNA library is thoughtfully presented in a tabular format, where sgRNAs are meticulously organized by their respective targeted genes and annotated with their corresponding start positions (see **Materials and Methods**). Additionally, an interactive page is generated using the D3GB genome browser [31], offering an intuitive visualization of the sgRNA library. This page accurately maps all genes and sgRNAs to their natural positions within the genome.

**Figure 2.**
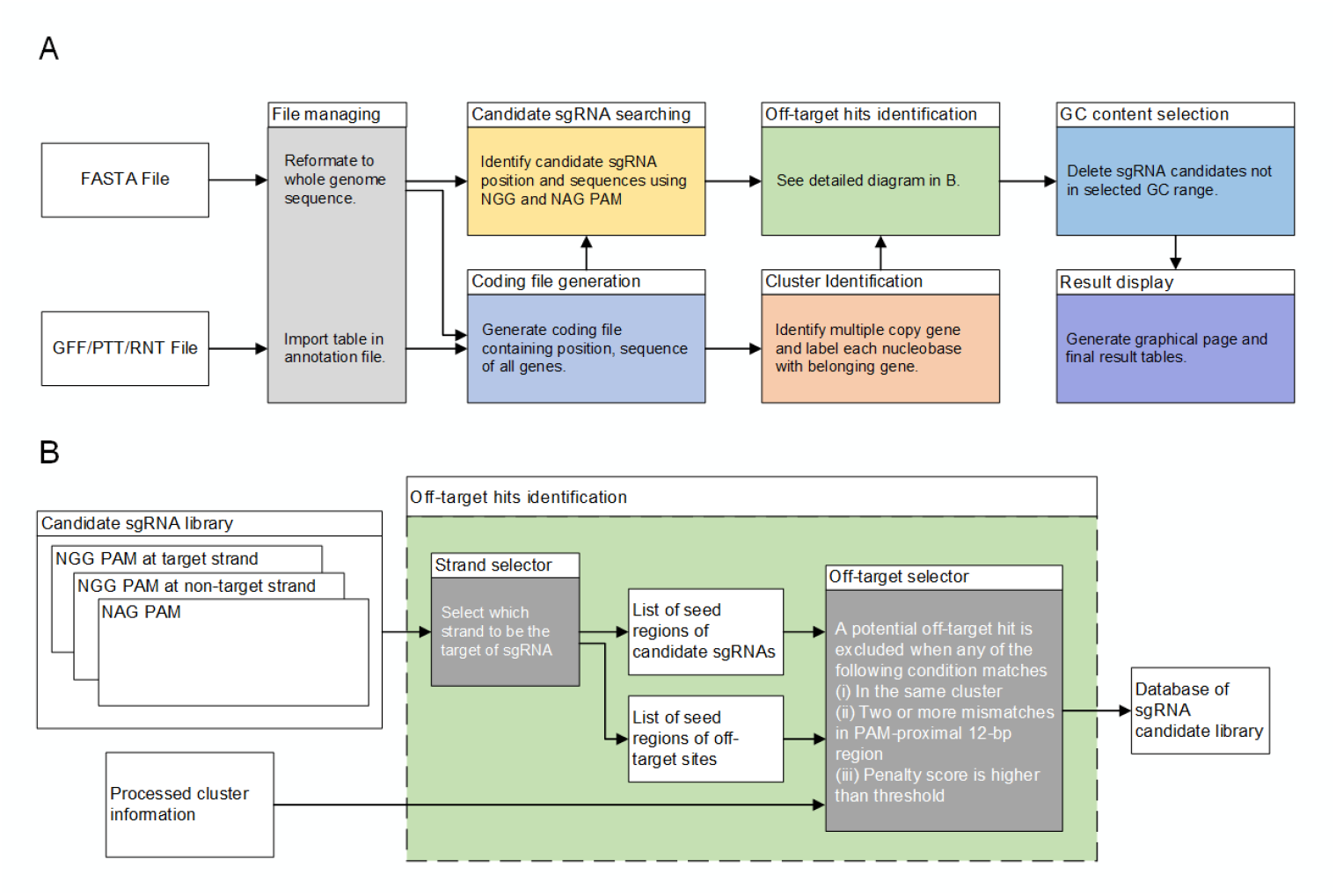
Functional blocks of GLiDe. **(A)** The GLiDe flowchart encompasses the processing of the FASTA file and the annotation file through multiple modules. These modules conduct candidate sgRNA searches, quality control (identifying off-target hits and selecting GC content), and ultimately present the results. **(B)** The off-target hits identification module generates three FASTA files containing 12 bp PAM-proximal sequences of sgRNA candidates. Alignments are performed among these three files to identify potential off-target hits.

### GLiDe is capable of designing CRISPRi sgRNA libraries

It is important to note that instead of CRISPR/Cas9 system that existing sgRNA design tools are focusing on, GLiDe is specially tailored for CRISPRi system. CRISPR/Cas9 system requires conformational gating process for formatting an R-loop structure before DNA cleavage [23], which requires sufficient base pairing between the sgRNA and the target DNA, reducing the likelihood of off-target cleavage. Consequently, sgRNAs without off-target cleavage may have potential off-target binding effects. GLiDe implements a more rigorous off-target selection principle, enabling the identification of these sgRNAs. To illustrate this capability, we compared sgRNA library of *Corynebacterium glutamicum* ATCC 13032 designed by GLiDe and sgRNA libraries generated by two widely used tools: CHOPCHOP [19] and Synthego (https://design.synthego.com/). We discovered that some of the highly ranked sgRNAs intended for CRISPR/Cas9 systems were indeed identified as having the potential for off-target binding. Among these identified off-target sequences, we randomly chose four (R1–R4, Table 1) for further validation.

**Table 1.**
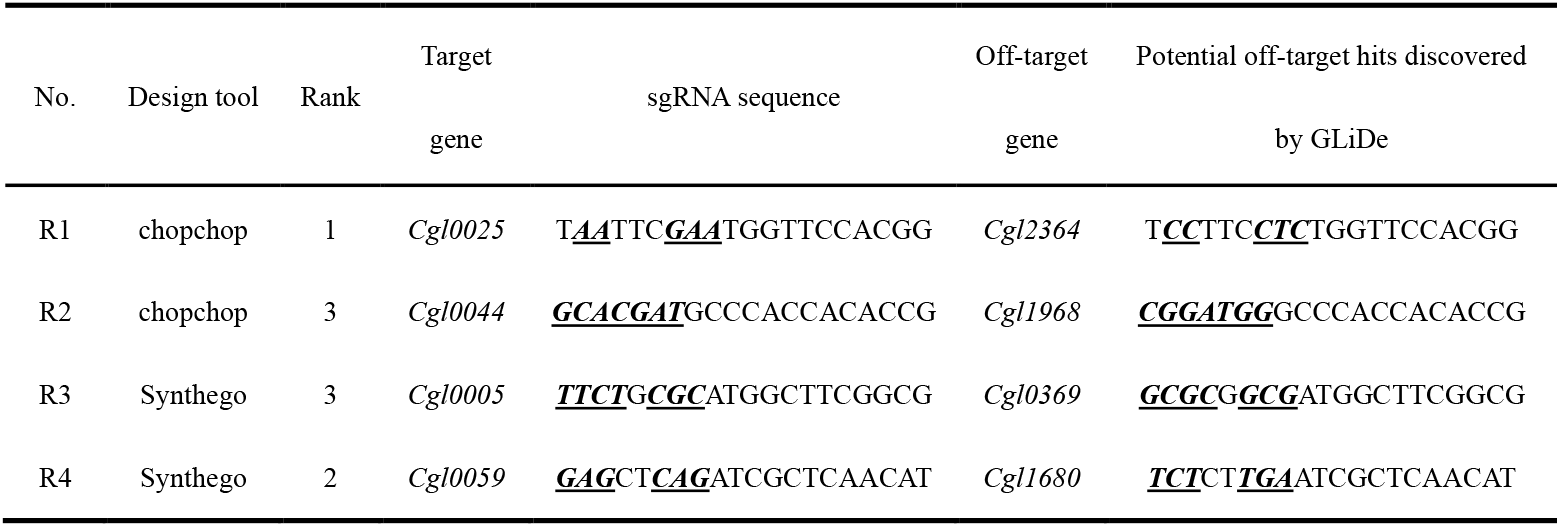
An example of four high rank sgRNAs designed by existing tools and their potential off-target hits identified by GLiDe.

To quantitatively assay the off-target binding effects, we applied a reporter system to connect binding affinity with mCherry expression [27]. In this system, we inserted the GLiDe identified off-target hit (N20 sequence) upstream of the *mcherry* promoter. Upon binding of the dCas9-sgRNA complex to the N20 sequence, the expression of *mcherry* would be repressed, resulting in altered fluorescence intensity. For each sgRNA (EXP) and its potential off-target hit, two additional sgRNAs were designed as positive (PC) and negative control (NC). The PC was the reverse-complement of the N20 sequence and was used to represent full repression; meanwhile, the NC had no binding affinity with the N20 sequence and was used to represent the fluorescence intensity without repression (Figure 3A). Since sgRNAs may bind to the endogenous genome to impact fluorescence intensity, experiments were conducted in *Escherichia coli* MCm, and all sgRNAs were cross-referenced with the bacterial genome to ensure the absence of potential off-target hits. Moreover, a blank control N20 sequence (blank) was designed, which had no binding affinity to any sgRNA, to assess whether the sgRNAs themselves had any influence on mCherry fluorescence. The sgRNA and N20 sequences used in all experiment groups are listed in Table 2.

**Table 2.**
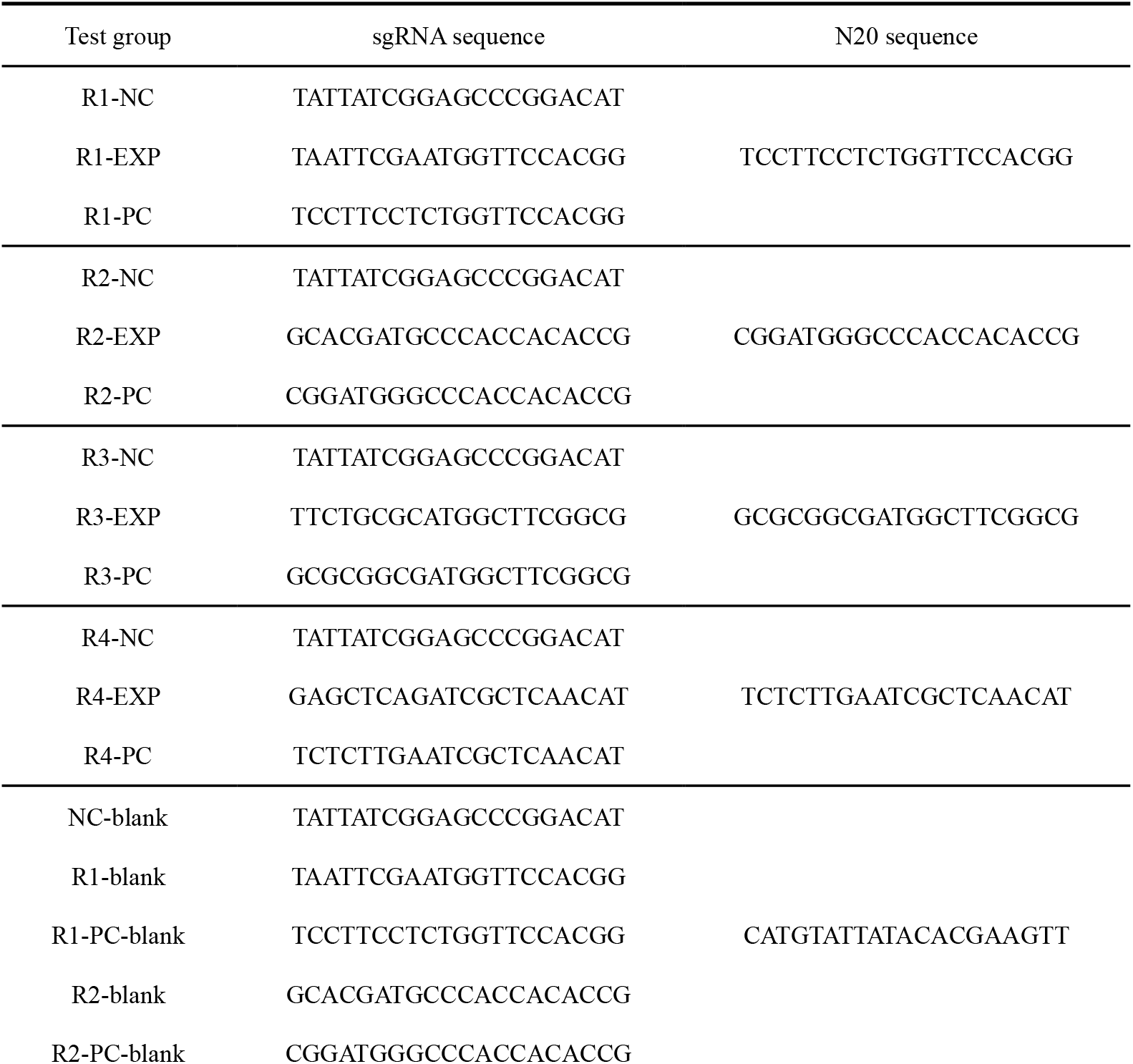
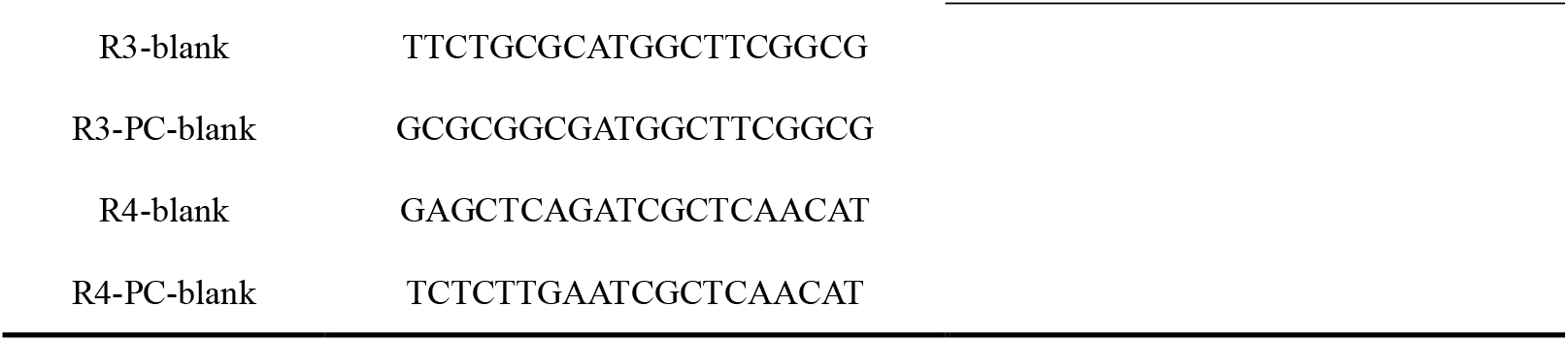
sgRNAs and N20 sequences of each test group.

**Figure 3.**
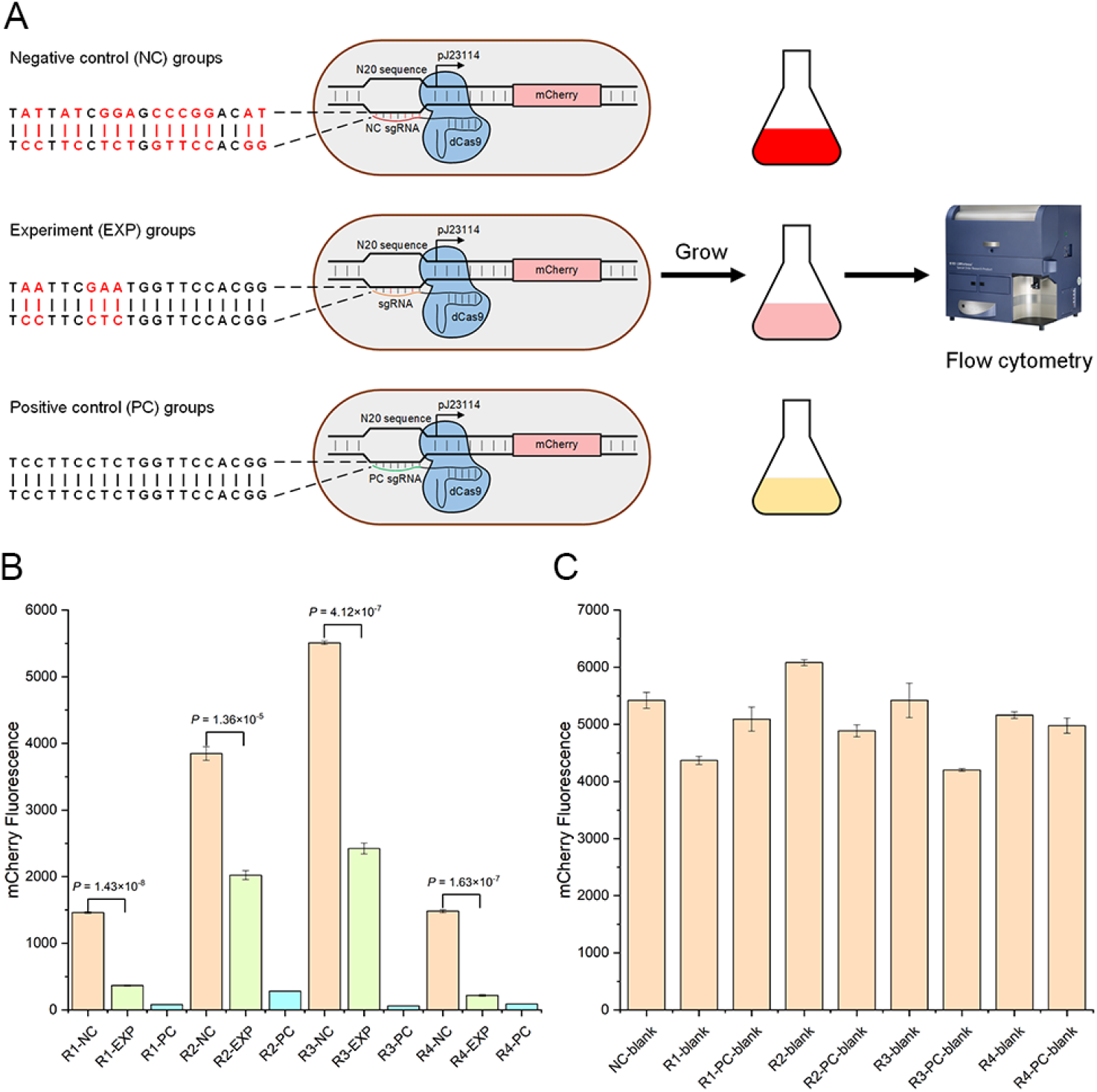
Assessment of off-target binding affinities using flow cytometry assay. **(A)** The binding affinity of each mismatched sgRNA was compared with positive controls and negative controls. Fluorescence intensities of the three groups were measured by flow cytometry. **(B)** All sgRNAs from the EXP groups exhibited significant off-target binding affinities (*P* < 10^−4^ for all groups, two tailed *t*-test). **(C)** The fluorescence intensities of the blank control groups were similar, demonstrated that the effects of sgRNA expression on *mcherry* expression are negligible. Therefore, the fluorescence intensity variations among EXP, PC, and NC groups were originated from the differences in binding affinities of sgRNAs. In both **(B)** and **(C)**, error bars represent the standard deviations of biological replicates (*n* = 3).

The results revealed that all four sgRNAs bound to the off-target hits identified by GLiDe, leading to reduced fluorescence intensity compared to the negative control groups (Figure 3B, *P* < 1.36 × 10^−5^ for all groups, two-tailed *t*-test; Figure S1). The repression rates ranged from 51.12% (R2) to 90.70% (R4), with an average of 69.47% (see **Materials and Methods**). This rate is capable of causing unexpected outcomes and potentially result in false positives in the screening experiment [11]. The blank control groups exhibited similar mCherry fluorescence intensities (Figure 3C), indicating that the observed repressions were primarily attributed to off-target effects. These findings provide strong evidence that GLiDe excels in the design of CRISPRi sgRNA libraries.

## DISCUSSION

GLiDe presents several distinctive advantages. Firstly, it concentrates on designing genome-scale CRISPRi sgRNA libraries, a focus notably lacking in prior web-based platforms like CHOPCHOP [19] and synthego (https://design.synthego.com/). Secondly, in contrast to existing scripts for bacterial CRISPRi sgRNA design [14,32,33], GLiDe provides a more convenient and flexible way for a wide range of user groups. As a web-based tool, the entire calculation process occurs online, eliminating the need for installation or local dependencies. Its compact, user-friendly interface simplifies the process, requiring researchers to set only a few parameters and optionally submit the sequence and annotation of the genome of interest. Finally, GLiDe’s quality control mechanism is built on a solid theoretical foundation. Two key design rules, the seed region rule (not yet considered by existing tools) and the penalty scoring rule, have been introduced based on extensive experimental evidence. This robust theoretical underpinning enhances the reliability of GLiDe, contributing to the accuracy and effectiveness of the tool. Our further experiments have demonstrated GLiDe’s superiority over existing tools when employed to design CRISPRi sgRNA libraries. Therefore, we believe GLiDe has extensive applicability in functional genomic studies such as essential genes study [34,35], bioprocesses optimization in industrially relevant microorganisms [36,37], or identification of complex genetic interactions [38].

The quality control methodology employed by GLiDe is based on the strand invasion model. During binding, mismatches in PAM-proximal region, particularly when there are two mismatches in seed region, could create a substantial energy barrier that cannot be adequately compensated. This barrier effectively impedes the strand invasion from the beginning, increasing the likelihood of dCas9 disengaging from the target. However, there is still potential limitation in the estimation of the impact of PAM-proximal mismatches. The current penalty scoring system primarily relies on experimental data, lacking a robust theoretical explanation. Furthermore, the current model assumes equal contributions from different mismatched nucleotide pairs, yet variations in stability exist among these mismatches [39]. Certain mismatches, such as rU/dG, rG/dT, and rG/dG, exhibit stabilities similar to Watson-Crick pairs [40]. One potential avenue for improvement is to incorporate thermodynamics. By treating the combination process as a Markov chain model, we can quantitatively calculate the thermodynamics of the sgRNA/DNA binding process based on fundamental thermodynamic parameters (nearest-neighbor parameters) [27]. We have further gathered nearest-neighbor parameters for the remaining 12 RNA/DNA single internal mismatches [41] and incorporated these data into the quantitative CRISPRi design tool we previously introduced (https://www.thu-big.net/sgRNA_design/Quantitative_CRISPRi_Design/). Nevertheless, the current tool faces limitations in computing binding activities when mismatches are adjacent or located at the ends due to a lack of corresponding thermodynamic parameters. The challenge arises because adjacent mismatches are too numerous to be directly measured by experiments. Substantial work is still needed to complete the thermodynamic data for these scenarios.

Additionally, in the current off-target hits identification process of GLiDe, hits in different locations are treated equally. However, it has been observed that the knock-down efficiency decreases when the sgRNA binding site is located far from the TSS [9]. Therefore, there is a possibility of ruling out sgRNAs that may not significantly impact downstream CRISPRi screening experiments. Despite this, the impact is considered acceptable due to the ample size of the designed library for CRISPRi selection. In addition, the optimal window for active sgRNA positioning relative to the transcription start site in a CRISPRi system varies across different organisms [42,43]. Consequently, the development of a unified approach for evaluating downstream effects becomes challenging due to these organism-specific differences. Overall, owing to the usability, solid basis, and high precision, we believe GLiDe can serve as a powerful tool that provides a wide range of novel research opportunities.

## MATERIALS AND METHODS

### Data prerequisites for the design of sgRNA library

To Design an sgRNA library, GLiDe requires two types of files: a FASTA file containing the entire DNA sequence, and an annotation file offering comprehensive details including genomic names, coding types, locations, strands, phases, and functional attributes. Regarding the annotation file, two formats are acceptable: (i) generic feature format files (GFF) and (ii) protein/RNA table files (PTT/RNT). These files can be easily obtained from a variety of publicly available databases such as NCBI [44], EMBL [45], DDBJ [46], etc. In the case of a newly sequenced organism, they can also be generated using genome annotation pipeline like PGAP [47].

### sgRNA library design workflow

As shown in Figure 2A, upon receiving the files, GLiDe constructs a coding list that connect each gene feature with its corresponding sequence. Genes with high similarity are considered as the same function [48,49] and are grouped into a cluster. This is achieved by employing BLASTN [50] with default parameters for sequence alignment, applying a strict threshold (< 0.001 evalue, > 95% identity, > 95% hit coverage, and > 95% query coverage). Each cluster may encompass one or more genes, and genes within the same cluster are considered to be multiple copies of the same gene. Consequently, potential hits across the cluster are not considered as off-target. For example, Table S1 provides a list of all clusters of multi-copy coding genes in *E. coli* MG1655. Candidate sgRNAs are identified using regular expressions, targeting two categories of guides: N20NGG and N20NAG (N= A, T, C or G), corresponding to canonical and non-canonical PAM sequences [9]. These candidate sgRNAs are categorized into three groups: (i) sgRNAs with an NGG PAM targeting the non-template strand; (ii) sgRNAs with an NGG PAM targeting the template strand; and (iii) sgRNAs with an NAG PAM. Three FASTA files (FASTA1, FASTA2, and FASTA3) are generated, corresponding to group (i), (ii), and (iii), respectively. These files contain the PAM-proximal 12-bp regions (seed regions) of all candidate sgRNAs, as these regions are pivotal for binding [14]. An illustrative example is presented in Figure S2. Following this, candidate sgRNAs undergo a rigorous quality control process to eliminate those with potential off-target hits (see “Quality control of sgRNA library” section). Subsequently, only sgRNAs adhering to the user-specified upper and lower limits for GC content are retained. Finally, the designed library is presented to the user, as detailed in the “Visualization of results” section.

### Quality control of sgRNA library

To improve the accuracy of off-target hit identification, our approach relies on the strand invasion model, which is grounded in the natural binding process [51–53] and extensive experimental data [8,54]. The fundamental concept involves identifying sequences that are less likely to result in off-target hits, with a focus on those exhibiting fewer mismatches in the PAM-proximal region compared to the target sequence. This process involves three alignments according to the user defined target strand (template or non-template) using SeqMap [55]. For instance, if the non-template strand is selected, FASTA1 would be aligned with FASTA1, FASTA2, and FASTA3. We implement two general rules to evaluate the impact of each off-target hit (Figure 2B).

One is seed region rule, where the seed region refers to the PAM-proximal 7–12 bp region. Mismatches within the seed region exert a notable impact on the binding affinity [14,56]. Our previous study revealed that two mismatches in the seed region substantially weaken binding affinity [27]. Despite its effectiveness, this principle has not been integrated into existing tools. Consequently, in our design, off-target binding is not deemed to occur when there are more than two mismatches within the 12 bp PAM-proximal region.

The second rule is the penalty scoring rule, which considers the influence of mismatches based on their distance to the PAM, considering that mismatches are generally better tolerated at the 5’ end of the sgRNA than at the 3’ end [18]. The sgRNA regions are categorized from the 3’ end to the 5’ end as region I (7 nt), region II (5 nt), and region III (the remaining sgRNA sequence). This division is informed by the experimental findings [15,24,57]. The mismatch penalties for region I, II, and III are 8, 4.5, and 2.5 (for NGG PAM), and 10, 7, and 3 (for NAG PAM), respectively (Figure S3). The effectiveness of this strategy has been demonstrated in a previous study [10]. An off-target site is identified when the penalty score falls below the user-defined threshold, with a recommended range of 18–21 for designing genome-scale libraries.

### Design of negative control sgRNAs

Negative control sgRNAs are designed to have no specific targets across the genome, which can be used to assess the influence of external factors on cellular phenotype. GLiDe designs these sgRNAs by generating random N20 sequences and subsequently removing those with notable target sites. For both NGG and NAG PAMs, a penalty score of 25 is applied, and the GC content limits match those of the sgRNA library. Additionally, GLiDe ensures that there are no five or more consecutive identical bases in these negative control sgRNAs.

### Visualization of results

GLiDe provides the final sgRNA library in a table, where each sgRNA is associated with its target gene and ranked according to its proximity to the start codon. This ranking is because sgRNAs targeting closer to transcription start site (TSS) have shown higher knock-down activity [9]. In addition to the table, GLiDe also generates an interactive graphical interface using D3GB [31]. This interface provides users with a comprehensive overview of the entire genome and the designed sgRNA library.

### DNA manipulation and reagents

Plasmid extraction and DNA purification procedures were carried out employing kits provided by Vazyme. PCR reactions were performed utilizing KOD One™ PCR Master Mix from TOYBO Life Science. The PCR primers were ordered from Azenta (Table S2). Plasmids were constructed by Gibson Assembly, with a mixture comprising 10 U/μL T5 exonuclease, 2 U/μL Phusion High-Fidelity DNA polymerase, and 40 U/μL Taq DNA ligase, all sourced from New England Biolabs. The antibiotic concentrations for kanamycin and ampicillin were maintained at 50 and 100 mg/L, respectively. A comprehensive list of all strains and plasmids utilized in this study can be found in Table S3.

### Plasmid construction

The reporting system was established using two separate plasmids: one harbored dCas9, named pdCas9-J23111, was previously constructed as described in prior work [10]. The other plasmid was derived from pN20test-114mCherry-r0-m1 [27], which was responsible for the expression of sgRNA and *mcherry*. In this particular plasmid, sgRNA expression was regulated by the J23119 promoter, while the N20 sequence was inserted upstream of the −35 region of the J23114 promoter, controlling *mcherry* expression. A total of 21 distinct pN20test-114mCherry plasmids were constructed (Figure S4). The plasmids were assembled using the Gibson Assembly method from PCR products (primers listed in Table S2), using the original pN20test-114mCherry-r0-m1 plasmid as the template. All constructed plasmids were confirmed through Sanger sequencing.

### Cell cultivation

Strains were initially cultured overnight at 37°C and 220 rpm in a 48-well deep-well plate, each contained 1 mL of LB medium with kanamycin and ampicillin. The grown cells were transferred to fresh LB medium with a 0.5% dilution and grown again under the same conditions as above for 10 hours. This subculture process was repeated to ensure the stability of mCherry expression and avoid cell adhesion. In preparation for cytometry assays, cultures were next diluted in fresh LB medium with antibiotics to OD600 = 0.02 and then grown for 4 hours to the logarithmic phase. After cultivation, 5 μL of culture medium from each well was diluted into 200 μL of phosphate-buffered saline. Three independent biological replicates were prepared for each strain.

### Flow cytometry assay and data processing

The flow cytometry assay was performed on an LSRFortessa flow cytometer (BD Biosciences) using a 96-well plate. Gating based on the FSC area and SSC area was carried out to exclude non-cell particles. To ensure accurate measurements, autofluorescence was quantified using the MCm/PdCas9-J23111 strain and subsequently subtracted during the data analysis process.

In the cytometry analysis, the fluorescence intensity distribution was log10-transformed and fitted to a two-component Gaussian mixture model [58] with parameters (*λ, μ*_1_, *μ*_2_, *σ*_1_, *σ*_2_) through the expectation-maximization algorithm. Here, *λ* and 1 − *λ* represent the mixing coefficients of the two Gaussian components, *μ*_1_, *μ*_2_, *σ*_1_ and *σ*_2_ represent the mean and standard deviation of the first and second Gaussian component, respectively (Equation 1).

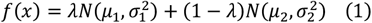

The mean expression strength was calculated with Equation 2.

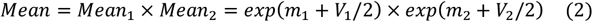

where 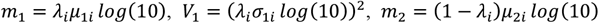 and 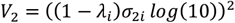.

The repression rate of each group was calculated using Equation 3.

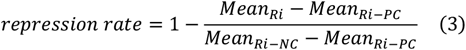

where *i* = 1–4.

## ACKNOWLEDGEMENT

We thank Dr. T. Wang (Tsinghua University) for building the original demo of the sgRNA design tool. We also thank members from the C. Zhang and H. Xing laboratories for critical discussions of this work.

## DATA AVAILABILITY

Raw data of the flow cytometry assays and plasmid maps related to this work can be accessed via GitHub (https://github.com/falconxtj/GLiDe).

## REFERENCES

1. van Opijnen T, Bodi KL, Camilli A. Tn-seq: high-throughput parallel sequencing for fitness and genetic interaction studies in microorganisms. Nat Methods 2009; 6:767–772

2. Garst AD, Bassalo MC, Pines G, et al. Genome-wide mapping of mutations at single-nucleotide resolution for protein, metabolic and genome engineering. Nat Biotechnol 2017; 35:48–55

3. Freed EF, Winkler JD, Weiss SJ, et al. Genome-Wide Tuning of Protein Expression Levels to Rapidly Engineer Microbial Traits. ACS Synth Biol 2015; 4:1244–1253

4. Feng H, Yuan Y, Yang Z, et al. Genome-wide genotype-phenotype associations in microbes. J Biosci Bioeng 2021; 132:1–8

5. Cong L, Ran FA, Cox D, et al. Multiplex genome engineering using CRISPR/Cas systems. Science (1979) 2013; 339:819–823

6. Mali P, Yang L, Esvelt KM, et al. RNA-Guided Human Genome Engineering via Cas9. Science (1979) 2013; 339:823–826

7. Bikard D, Jiang W, Samai P, et al. Programmable repression and activation of bacterial gene expression using an engineered CRISPR-Cas system. Nucleic Acids Res 2013; 41:7429–7437

8. Jinek M, Chylinski K, Fonfara I, et al. A Programmable Dual-RNA–Guided DNA Endonuclease in Adaptive Bacterial Immunity. Science (1979) 2012; 337:816–821

9. Qi LS, Larson MH, Gilbert LA, et al. Repurposing CRISPR as an RNA-guided platform for sequence-specific control of gene expression. Cell 2013; 152:1173–1183

10. Wang T, Guan C, Guo J, et al. Pooled CRISPR interference screening enables genome-scale functional genomics study in bacteria with superior performance. Nat Commun 2018; 9:2475

11. Hawkins JS, Silvis MR, Koo BM, et al. Mismatch-CRISPRi Reveals the Co-varying Expression-Fitness Relationships of Essential Genes in Escherichia coli and Bacillus subtilis. Cell Syst 2020; 11:523–535.e9

12. Lian J, Schultz C, Cao M, et al. Multi-functional genome-wide CRISPR system for high throughput genotype– phenotype mapping. Nature Communications 2019 10:1 2019; 10:1–10

13. Chen P, Michel AH, Zhang J. Transposon insertional mutagenesis of diverse yeast strains suggests coordinated gene essentiality polymorphisms. Nature Communications 2022 13:1 2022; 13:1–15

14. de Bakker V, Liu X, Bravo AM, et al. CRISPRi-seq for genome-wide fitness quantification in bacteria. Nature Protocols 2021 17:2 2022; 17:252–281

15. Hsu PD, Scott DA, Weinstein JA, et al. DNA targeting specificity of RNA-guided Cas9 nucleases. Nat Biotechnol 2013; 31:827–832

16. Shalem O, Sanjana NE, Zhang F. High-throughput functional genomics using CRISPR–Cas9. Nat Rev Genet 2015; 16:299–311

17. Heigwer F, Kerr G, Boutros M. E-CRISP: fast CRISPR target site identification. Nat Methods 2014; 11:122–1. 123

18. Sander JD, Joung JK. CRISPR-Cas systems for editing, regulating and targeting genomes. Nat Biotechnol 2014; 32:347–355

19. Montague TG, Cruz JM, Gagnon JA, et al. CHOPCHOP: a CRISPR/Cas9 and TALEN web tool for genome editing. Nucleic Acids Res 2014; 42:W401–W407

20. Concordet J-P, Haeussler M. CRISPOR: intuitive guide selection for CRISPR/Cas9 genome editing experiments and screens. Nucleic Acids Res 2018; 46:W242–W245

21. Bae S, Park J, Kim J-S. Cas-OFFinder: a fast and versatile algorithm that searches for potential off-target sites of Cas9 RNA-guided endonucleases. Bioinformatics 2014; 30:1473–1475

22. Liu H, Wei Z, Dominguez A, et al. CRISPR-ERA: a comprehensive design tool for CRISPR-mediated gene editing, repression and activation: Fig. 1. Bioinformatics 2015; 31:3676–3678

23. Sternberg SH, LaFrance B, Kaplan M, et al. Conformational control of DNA target cleavage by CRISPR– Cas9. Nature 2015; 527:110–113

24. Boyle EA, Andreasson JOL, Chircus LM, et al. High-throughput biochemical profiling reveals sequence determinants of dCas9 off-target binding and unbinding. Proceedings of the National Academy of Sciences 2017; 114:5461–5466

25. Struhl K. Fundamentally Different Logic of Gene Regulation in Eukaryotes and Prokaryotes. Cell 1999; 98:1– 4

26. Kuzminov A. The Precarious Prokaryotic Chromosome. J Bacteriol 2014; 196:1793–1806

27. Feng H, Guo J, Wang T, et al. Guide-target mismatch effects on dCas9–sgRNA binding activity in living bacterial cells. Nucleic Acids Res 2021; 49:1263–1277

28. Gaudelli NM, Komor AC, Rees HA, et al. Programmable base editing of A•T to G•C in genomic DNA without DNA cleavage. Nature 2017; 551:464–471

29. Komor AC, Kim YB, Packer MS, et al. Programmable editing of a target base in genomic DNA without double-stranded DNA cleavage. Nature 2015 533:7603 2016; 533:420–424

30. Chen B, Gilbert LA, Cimini BA, et al. Dynamic Imaging of Genomic Loci in Living Human Cells by an Optimized CRISPR/Cas System. Cell 2013; 155:1479–1491

31. Barrios D, Prieto C. D3GB: An Interactive Genome Browser for R, Python, and WordPress. J Comput Biol 2017; 24:447–449

32. Liu X, Kimmey JM, Matarazzo L, et al. Exploration of Bacterial Bottlenecks and Streptococcus pneumoniae Pathogenesis by CRISPRi-Seq. Cell Host Microbe 2021; 29:107–120.e6

33. Banta AB, Ward RD, Tran JS, et al. Programmable Gene Knockdown in Diverse Bacteria Using Mobile-CRISPRi. Curr Protoc Microbiol 2020; 59:e130

34. Peters JM, Colavin A, Shi H, et al. A Comprehensive, CRISPR-based Functional Analysis of Essential Genes in Bacteria. Cell 2016; 165:1493–1506

35. Rousset F, Cabezas-Caballero J, Piastra-Facon F, et al. The impact of genetic diversity on gene essentiality within the Escherichia coli species. Nat Microbiol 2021; 6:301–312

36. Li S, Jendresen CB, Landberg J, et al. Genome-Wide CRISPRi-Based Identification of Targets for Decoupling Growth from Production. ACS Synth Biol 2020; 9:1030–1040

37. Donati S, Kuntz M, Pahl V, et al. Multi-omics Analysis of CRISPRi-Knockdowns Identifies Mechanisms that Buffer Decreases of Enzymes in E. coli Metabolism. Cell Syst 2021; 12:56–67.e6

38. Jaffe M, Dziulko A, Smith JD, et al. Improved discovery of genetic interactions using CRISPRiSeq across multiple environments. Genome Res 2019; 29:668–681

39. Sugimoto N, Nakano S, Katoh M, et al. Thermodynamic Parameters To Predict Stability of RNA/DNA Hybrid 1. Duplexes. Biochemistry 1995; 34:11211–11216

40. Sugimoto N, Nakano M, Nakano S. Thermodynamics−Structure Relationship of Single Mismatches in RNA/DNA Duplexes. Biochemistry 2000; 39:11270–11281

41. Xiang T, Feng H, Xing X, et al. Thermodynamic Parameters Contributions of Single Internal Mismatches In RNA/DNA Hybrid Duplexes. bioRxiv 2022; 2022.11.25.517909

42. Guo J, Wang T, Guan C, et al. Improved sgRNA design in bacteria via genome-wide activity profiling. Nucleic Acids Res 2018; 46:7052–7069

43. Smith JD, Suresh S, Schlecht U, et al. Quantitative CRISPR interference screens in yeast identify chemical-genetic interactions and new rules for guide RNA design. Genome Biol 2016; 17:1–16

44. Sayers EW, Bolton EE, Brister JR, et al. Database resources of the National Center for Biotechnology Information in 2023. Nucleic Acids Res 2023; 51:D29–D38

45. Thakur M, Bateman A, Brooksbank C, et al. EMBL’s European Bioinformatics Institute (EMBL-EBI) in 2022. Nucleic Acids Res 2023; 51:D9–D17

46. Tanizawa Y, Fujisawa T, Kodama Y, et al. DNA Data Bank of Japan (DDBJ) update report 2022. Nucleic Acids Res 2023; 51:D101–D105

47. Li W, O’Neill KR, Haft DH, et al. RefSeq: expanding the Prokaryotic Genome Annotation Pipeline reach with protein family model curation. Nucleic Acids Res 2021; 49:D1020–D1028

48. McTavish H, LaQuier F, Arciero D, et al. Multiple copies of genes coding for electron transport proteins in the bacterium Nitrosomonas europaea. J Bacteriol 1993; 175:2445–2447

49. Schrider DR, Hahn MW. Gene copy-number polymorphism in nature. Proceedings of the Royal Society B: Biological Sciences 2010; 277:3213–3221

50. Johnson M, Zaretskaya I, Raytselis Y, et al. NCBI BLAST: a better web interface. Nucleic Acids Res 2008; 36:W5–W9

51. Szczelkun MD, Tikhomirova MS, Sinkunas T, et al. Direct observation of R-loop formation by single RNA-guided Cas9 and Cascade effector complexes. Proc Natl Acad Sci U S A 2014; 111:9798–9803

52. Nishimasu H, Ran FA, Hsu PD, et al. Crystal Structure of Cas9 in Complex with Guide RNA and Target DNA. Cell 2014; 156:935–949

53. Cofsky JC, Soczek KM, Knott GJ, et al. CRISPR–Cas9 bends and twists DNA to read its sequence. Nat Struct Mol Biol 2022; 29:395–402

54. Jones SK, Hawkins JA, Johnson N V., et al. Massively parallel kinetic profiling of natural and engineered CRISPR nucleases. Nature Biotechnology 2020 39:1 2020; 39:84–93

55. Jiang H, Wong WH. SeqMap: mapping massive amount of oligonucleotides to the genome. Bioinformatics 2008; 24:2395–2396

56. Semenova E, Jore MM, Datsenko KA, et al. Interference by clustered regularly interspaced short palindromic repeat (CRISPR) RNA is governed by a seed sequence. Proc Natl Acad Sci U S A 2011; 108:10098–10103

57. Gilbert LA, Horlbeck MA, Adamson B, et al. Genome-Scale CRISPR-Mediated Control of Gene Repression and Activation. Cell 2014; 159:647–661

58. Feng H, Li F, Wang T, et al. Deep-learning–assisted Sort-Seq enables high-throughput profiling of gene expression characteristics with high precision. Sci Adv 2023; 9:

