## Supplementary Materials for "GLiDe: a web-based genome-scale CRISPRi sgRNA design tool for prokaryotes"

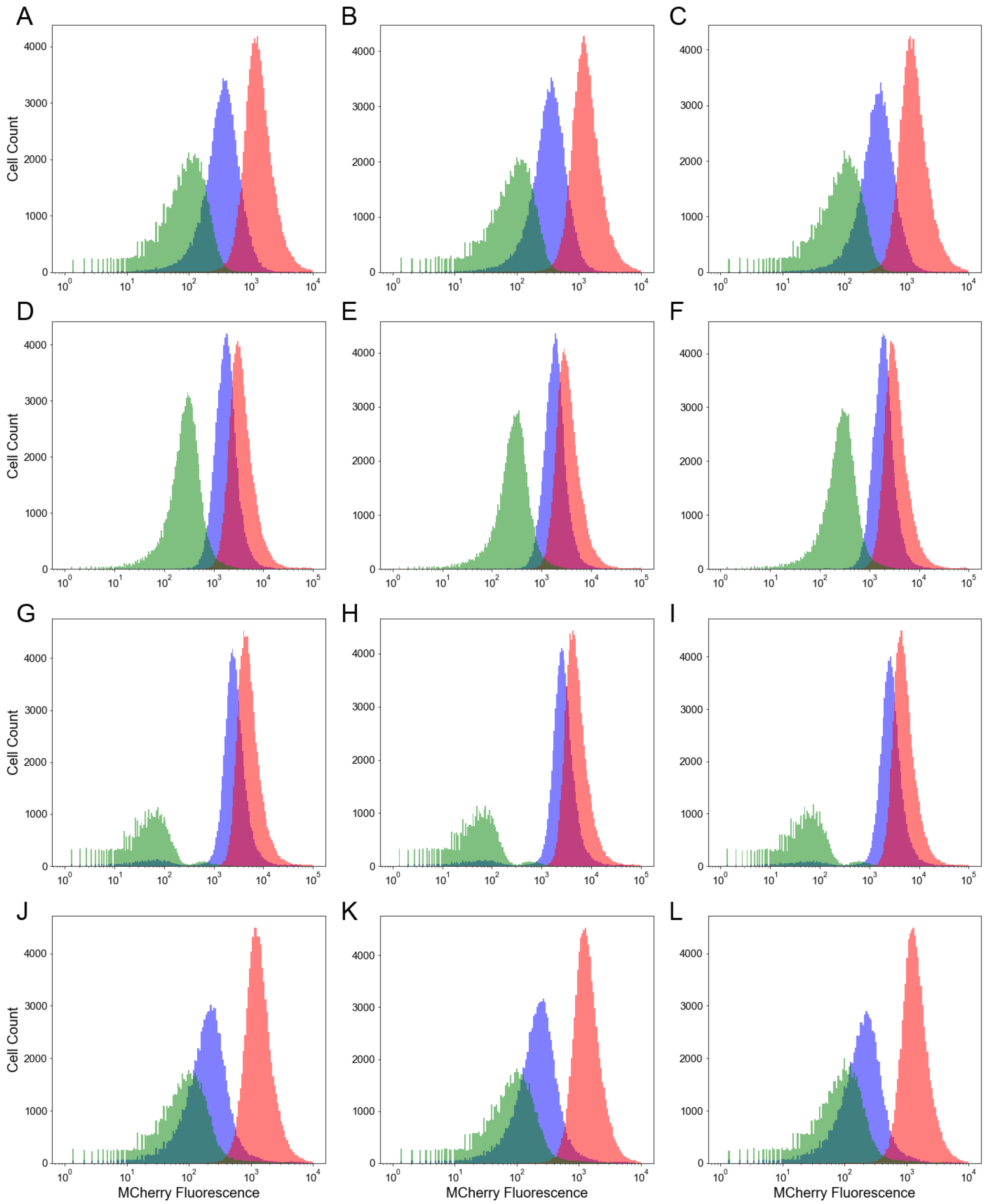


**Figure S1** Fluorescence intensity distributions of EXP (blue), NC (red) and PC groups (green). Three independent biological replicates were conducted for **(A-C)** R1, **(D-F)** R2, **(G-I)** R3, and **(J-L)** R4.

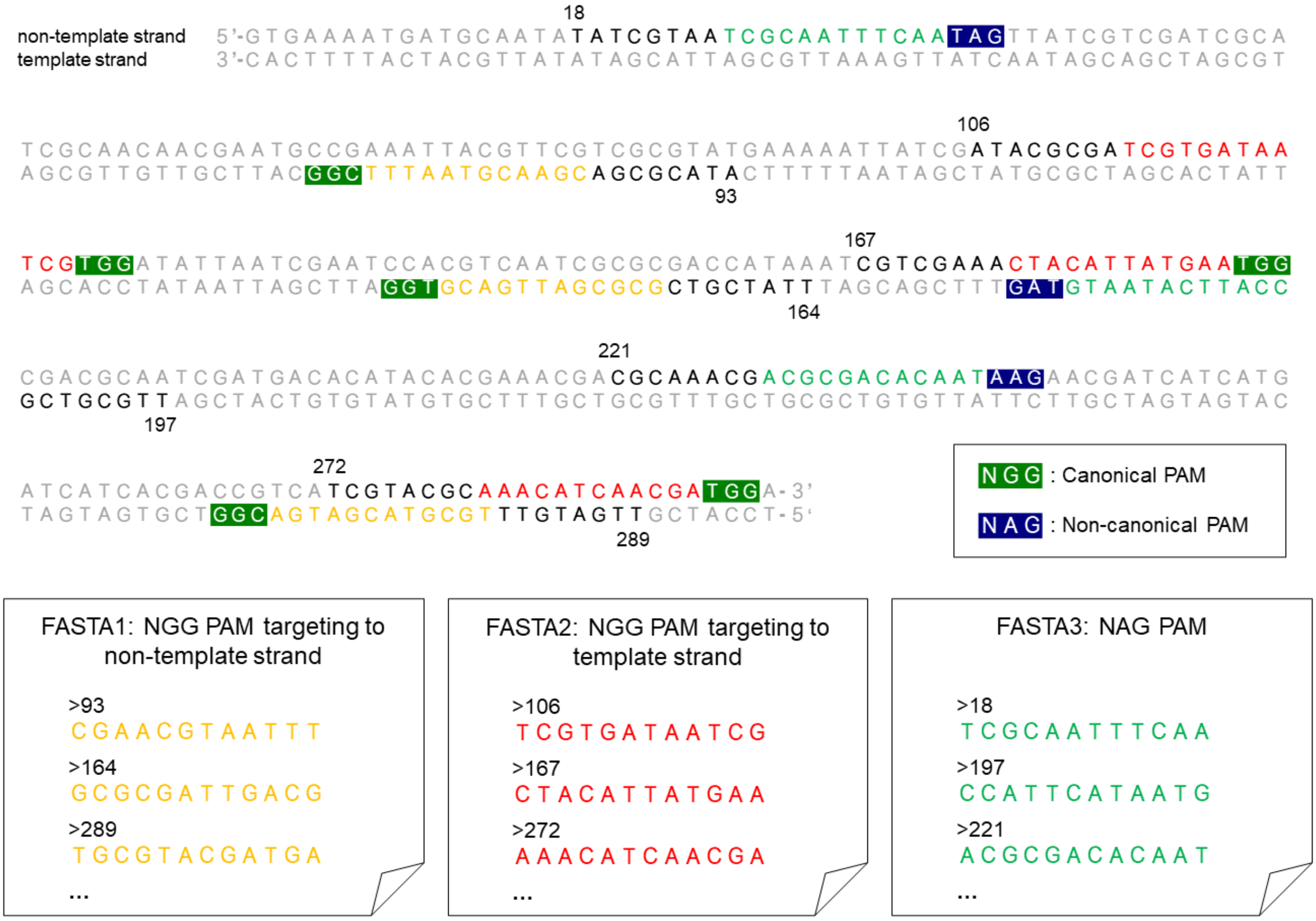


**Figure S2.** A toy example to illustrate the identification and classification of candidate sgRNAs. sgRNAs are searched by regular expressions and classified into three groups based on the PAM and targeted strands. The start position and PAM-proximal 12-bp of each sgRNA are saved in three separate FASTA files.

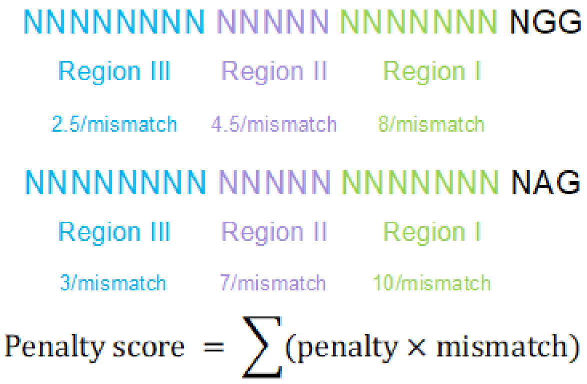


**Figure S3.** Schematic diagram of the calculation of penalty scores

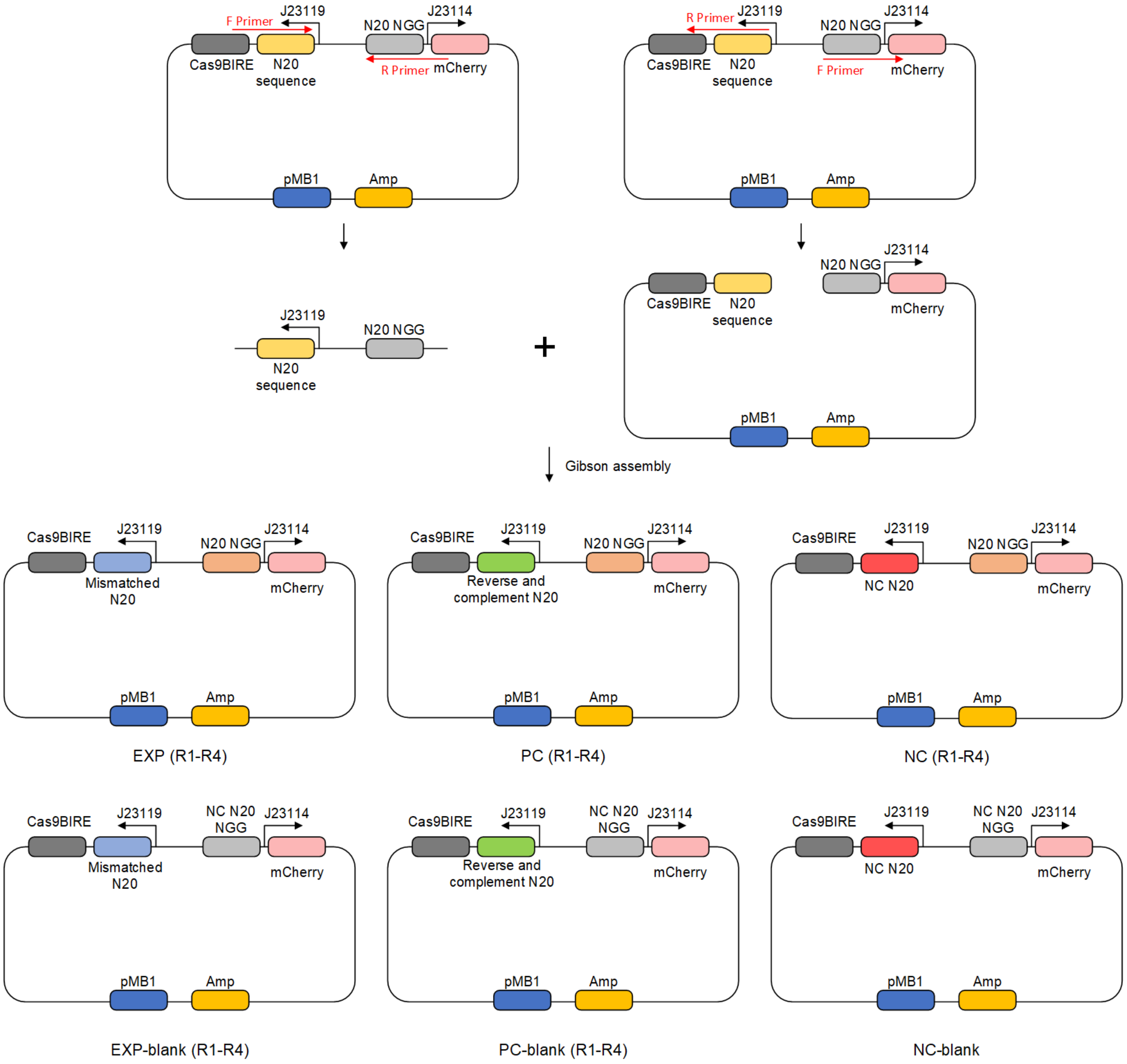


**Figure S4.** Workflow of the plasmid construction. The sgRNA and N20 sequences were introduced via Gibson assembly.

**Table S1:** Clusters of all multi-copy coding genes of *E. coli* MG1655 identified by GLiDe

| Gene name | Copy number |
|---|---|
| *insL* | 3 |
| *insB* | 7 |
| *insA* | 7 |
| *insI* | 3 |
| *insH* | 11 |
| *insE* | 5 |
| *insF* | 5 |
| *insC* | 6 |
| *insD* | 6 |
| *rzpD, rzoR* | 2 |
| *ylcI, ynfO* | 2 |
| *nohD, nohA* | 2 |
| *rhsC, rhsB, rhsA* | 3 |
| *ybfD, ydcC, yhhI* | 3 |
| *ldrA, ldrB, ldrC* | 3 |
| *tpr, ychS* | 2 |
| *tfaR, tfaQ* | 2 |
| *pinR, pinQ* | 2 |
| *ynaE, ydfK* | 2 |
| *ynaM, ynfT* | 2 |
| *hokB, mokB* | 2 |
| *gadB, gadA* | 2 |
| *yqgG, yqgC* | 2 |
| *tufA, tufB* | 2 |
| *yriA, yriB* | 2 |
| *cyaY, yzcX* | 2 |

**Table S2:** PCR Primers used in this study

| Name | Sequence |
|---|---|
| R1_F | CCGTGGAACCATTCGAATTAACTAGTATTATACCTAGGACTGAGC |
| R1_R | CTAGTTAATTCGAATGGTTCCACGGGTTTTAGAGCTAGAAATAGCAAGTT |
| N20_R1_F | TCCTTCCTCTGGTTCCACGGCGGTTTATGGCTAGCTCAGTCCTAG |
| N20_R1_R | AACCGCCGTGGAACCAGAGGAAGGAGTTGGATGTACTGCGGCTCCGTCTA |
| R1_PC_F | CCGTGGAACCAGAGGAAGGAACTAGTATTATACCTAGGACTGAGC |
| R1_PC_R | CTAGTTCCTTCCTCTGGTTCCACGGGTTTTAGAGCTAGAAATAGCAAGTT |
| R2_F | CGGTGTGGTGGGCATCGTGCACTAGTATTATACCTAGGACTGAGC |
| R2_R | CTAGTGCACGATGCCCACCACACCGGTTTTAGAGCTAGAAATAGCAAGTT |
| N20_R2_F | CGGATGGGCCCACCACACCGCGGTTTATGGCTAGCTCAGTCCTAG |
| N20_R2_R | AACCGCGGTGTGGTGGGCCCATCCGGTTGGATGTACTGCGGCTCCGTCTA |
| R2_PC_F | CGGTGTGGTGGGCCCATCCGACTAGTATTATACCTAGGACTGAGC |
| R2_PC_R | CTAGTCGGATGGGCCCACCACACCGGTTTTAGAGCTAGAAATAGCAAGTT |
| R3_F | CGCCGAAGCCATGCGCAGAAACTAGTATTATACCTAGGACTGAGC |
| R3_R | CTAGTTTCTGCGCATGGCTTCGGCGGTTTTAGAGCTAGAAATAGCAAGTT |
| N20_R3_F | GCGCGGCGATGGCTTCGGCGCGGTTTATGGCTAGCTCAGTCCTAG |
| N20_R3_R | AACCGCGCCGAAGCCATCGCCGCGCGTTGGATGTACTGCGGCTCCGTCTA |
| R3_PC_F | CGCCGAAGCCATCGCCGCGCACTAGTATTATACCTAGGACTGAGC |
| R3_PC_R | CTAGTGCGCGGCGATGGCTTCGGCGGTTTTAGAGCTAGAAATAGCAAGTT |
| R4_F | ATGTTGAGCGATCTGAGCTCACTAGTATTATACCTAGGACTGAGC |
| R4_R | CTAGTGAGCTCAGATCGCTCAACATGTTTTAGAGCTAGAAATAGCAAGTT |
| N20_R4_F | TCTCTTGAATCGCTCAACATCGGTTTATGGCTAGCTCAGTCCTAG |
| N20_R4_R | AACCGATGTTGAGCGATTCAAGAGAGTTGGATGTACTGCGGCTCCGTCTA |
| R4_PC_F | ATGTTGAGCGATTCAAGAGAACTAGTATTATACCTAGGACTGAGC |
| R4_PC_R | CTAGTTCTCTTGAATCGCTCAACATGTTTTAGAGCTAGAAATAGCAAGTT |
| NC_F | ATGTCCGGGCTCCGATAATAACTAGTATTATACCTAGGACTGAGC |
| NC_R | CTAGTTATTATCGGAGCCCGGACATGTTTTAGAGCTAGAAATAGCAAGTT |
| N20_NC_F | CCAACCATGTATTATACACGAAGTTCGGTTTATGGCTAGCTCAGTCCTAG |
| N20_NC_R | AACTTCGTGTATAATACATGGTTGGATGTACTGCGGCTCCGTCTA |

**Table S3:** Strains and plasmids used in this study

| Strains/plasmids | Sequence | Sources |
|---|---|---|
| **Strains** | | |
| *E. coli* K12 MG1655 | Wild type | ATCC 700936 |
| *E. coli* MCm | Carries chloromycetin expression cassette integrated into *smf* locus of *E. coli* K12 MG1655 | Ref (1) |
| **Plasmids** | | |
| pdCas9-J23111 | Expresses dCas9, p15A, $Km^R$ | Ref (2) |
| pN20test-J23114-MCherry-R1-EXP<br>pN20test-J23114-MCherry-R1-PC<br>pN20test-J23114-MCherry-R1-NC<br>pN20test-J23114-MCherry-R2-EXP<br>pN20test-J23114-MCherry-R2-PC<br>pN20test-J23114-MCherry-R2-NC<br>pN20test-J23114-MCherry-R3-EXP<br>pN20test-J23114-MCherry-R3-PC<br>pN20test-J23114-MCherry-R3-NC<br>pN20test-J23114-MCherry-R4-EXP<br>pN20test-J23114-MCherry-R4-PC<br>pN20test-J23114-MCherry-R4-NC | Express MCherry with different exp/NC/PC sgRNA and N20 sequence, pMB1, $Amp^R$ | This study |
| pN20test-J23114-MCherry-R1-EXP-blank<br>pN20test-J23114-MCherry-R1-PC-blank<br>pN20test-J23114-MCherry-R2-EXP-blank<br>pN20test-J23114-MCherry-R2-PC-blank<br>pN20test-J23114-MCherry-R3-EXP-blank<br>pN20test-J23114-MCherry-R3-PC-blank<br>pN20test-J23114-MCherry-R4-EXP-blank<br>pN20test-J23114-MCherry-R4-PC-blank<br>pN20test-J23114-MCherry-NC-blank | Express MCherry with negative control N20 sequence and different exp/NC/PC sgRNA, pMB1, $Amp^R$ | This study |

**References**


1. Wang,T., Guan,C., Guo,J., Liu,B., Wu,Y., Xie,Z., Zhang,C. and Xing,X.-H. (2018) Pooled CRISPR interference screening enables genome-scale functional genomics study in bacteria with superior performance. *Nat Commun*, **9**, 2475.
2. Feng,H., Guo,J., Wang,T., Zhang,C. and Xing,X. (2021) Guide-target mismatch effects on dCas9–sgRNA binding activity in living bacterial cells. *Nucleic Acids Res*, **49**, 1263–1277.
